# Detection of HER2 expression and its structural alterations in gastric cancer tissues infected with cagA+ *H. pylori*

**DOI:** 10.1101/2020.03.05.979963

**Authors:** Akhileshwar Kumar Srivastava, Divya Singh

**Author notes:** The National Institute for Biotechnology in the Negev, Ben-Gurion University of the Negev, Beer-Sheva 84105 Israel (Akhileshwar Kumar Srivastava).

## Abstract

**Background:** *Helicobacter pylori* (HP) cagA is the causing agent for development of gastric cancer (GC). *H. pylori* also involves to trigger the EGFR (epidermal growth factor receptor) expression in gastric cancer cells. However, the prognostic relation of cagA with HER2 status in GC was not well understood.

**Objective:** The main aim of this study was to investigate the link of HER2 expression with CagA+ *H. pylori* in GC tissues.

**Materials and Methods:** The study was performed on 85 GC tissues of GC patients. The specific primers of 16S rDNA and cagA for PCR amplification were used. For investigation of HER2 status in GC tissues, immunohistochemistry and PCR amplification were performed. In silico study was performed for the investigation of interactive potential of HER2 with CagA protein.

**Results:** PCR amplified the 54 (63.52 %) of 85 GC tissues for HP that showed 34 (62.96 %) cagA+ HP. Immunohistochemistry of tissues revealed 57 (67.05 %) diffuse and 28 (32.94 %) intestinal type cancer. Of 85 cases, 21 GC tissues scored (2 + or 3 +) for positive HER2 expression and score (0 or 1 +) of 64 (75.29 %) showed negative. Of 21 HER2 + GC tissue, 15 biopsies had cagA+ HP and 2 were negative. PCR amplified single amplicon in 17 (20 %) CagA+ tissues and 3 (5.55 %) in CagA - HP. The molecular interactions of CagA was also showed its efficiency for HER2 expression.

**Conclusion:** The study concluded that CagA+ HP may induce HER2 overexpression in GC tissues.

## INTRODUCTION

About half the human world population are highly affected by persistent colonization of *Helicobacter pylori* in the human stomach. This bacterium is known as causing agent for chronic gastric inflammation, which can change into severe diseases like peptic ulcer, mucosa-associated lymphoid tissue (MALT) and after prolongation of diseases finally lead to gastric cancer^[1,2]^. Several virulence genes of *H. pylori* in have been recognised among which the cytotoxin-associated antigene gene (cagA) and vacuolating cytotoxin antigen gene (vacA) that play role in pathogenicity to an increased risk of developing gastric atrophy^[3]^.

The status of molecule HER2 (human epidermal growth factor receptor 2) expression plays as a key role in the development and progression of many cancers, but it has a poor prognosis. Sakai et al. (1986) described first about overexpression of HER2 protein in gastric cancer using IHC (Immunohistochemistry)^[4]^. The HER2 molecules was amplified at rate of 20 % in breast cancer^[5]^. whereas in gastric cancer this amplification or protein overexpression was varied (7-34 %)^[6]^. The expression of HER2 in the lymph has own indication of carcinogenic activity and participates in clonal proliferation due to possible intra-tumoral heterogeneity in a gastric cancer case, which metastasized to gastric lymph nodes. Lymph node expression should be defined in relation of HER2 expression in tumor cell which metastasized into gastric lymph node^[7]^.

An earlier study reported that *H. pylori* has played a major role in the expression of EGFR (epidermal growth factor receptor) in gastric mucosal cells and enhanced the growth, proliferation and differentiation in cells^[8]^. Several studies explained that the level of EGF and EGFR were decreased in cells after eradication of *H. pylori* from cancerous tissues^[8]^. These studies indicated that EGF and EGFR levels were increased in subjects with *H. pylori*, and eradication had the vital importance^[8]^. The limited studies were performed on gastric cancer patients with positive expression of HER2 by *H. pylori*. Significant research progress has not been made about any link a major virulence gene cagA of *H. pylori* with HER2 status in infected sites of gastric cancer. The aim of the present study to determine the association of cagA gene with HER2 status in GC patients infected with *H. pylori*.

## MATERIAL AND METHODS

### Study of patients

Eighty five patients underwent for surgery between January 2017 and October 2018 at Institute of Medical Sciences, Banaras Hindu University, India. The patients included 58 (68.23 %) male and 27 (31.76 %) female, with age ranging 25-75 years (mean age, 56.22 ± 8.91 years). The biopsies of gastric cancer and healthy controls (normal gastric mucosa) were collected from each patients with prior consent. All clinical characteristics of patients are summarised in Table (1). Tissue samples were obtained during resection surgery. The samples were divided into three parts, one was used for rapid urease test (RUT), the 2nd part immediately frozen in liquid nitrogen, and stored at −80 ^°^C for DNA extraction and PCR assay for detection of *H. pylori* as well as HER2 status and 3rd part for immunohistochemical analysis.

**Table 1.**
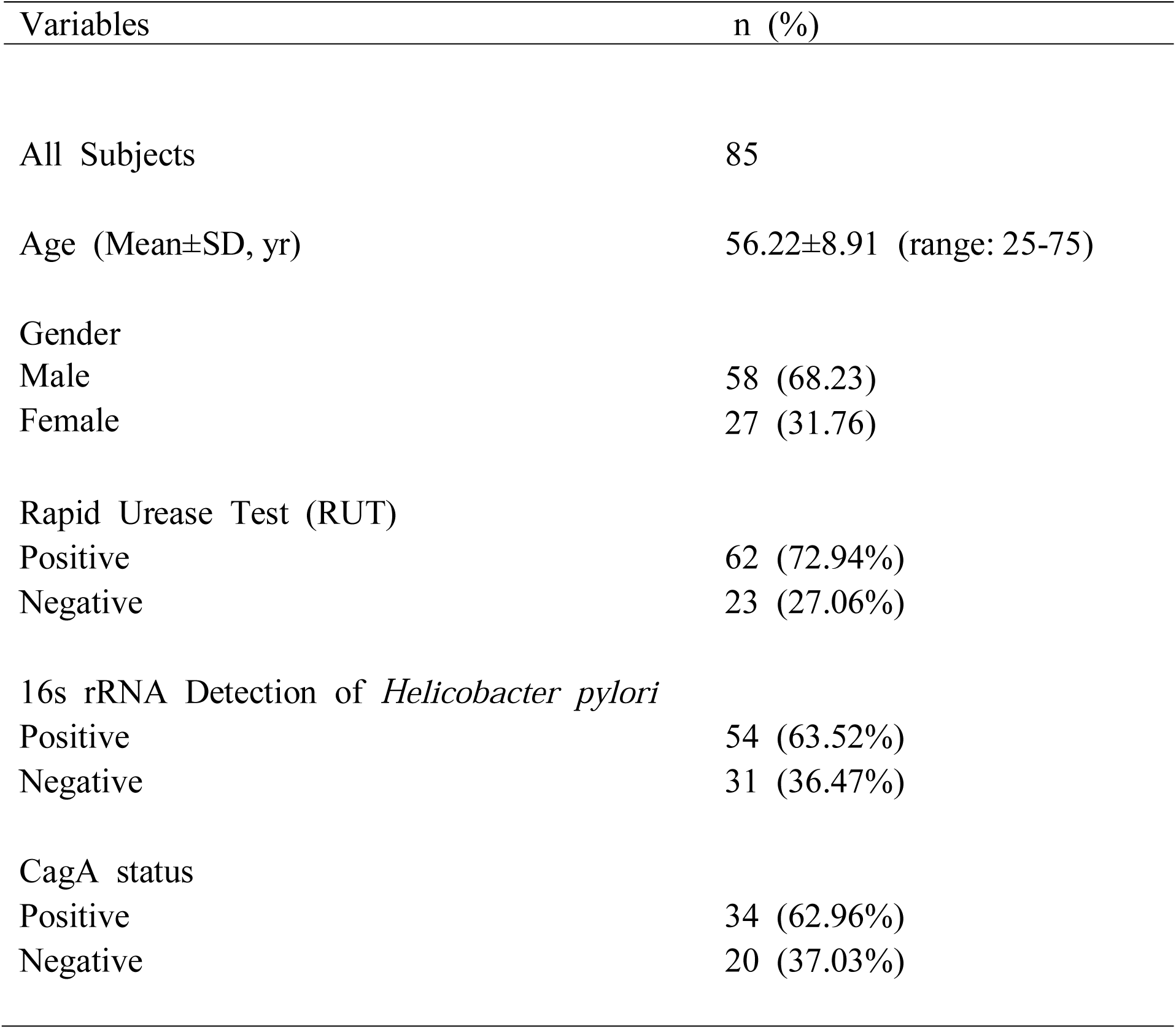
Characteristic of subjects.

### Bacterial strain

A strain, namely, *H. pylori* 26695 was used as reference isolate. The strain was positive for cagA gene. *H. pylori* 26695 was provided by Dr. Rukhsana Chaudhary, Indian Institute of Chemical Biology, Kolkata, India.

### Rapid urease test (RUT)

Rapid urease test was done, an indirect method for detection of presence of *H. pylori* in test as described by Thillainayagam (1991)^[9]^. The biopsy sample (1^st^) of each patients was kept for 1 h in sterile vial tube containing 1 ml of 10 % urea solution in deionized water (pH 6.8), and 2 drops of 1 % phenol red. A positive result was recorded by a change in the color of solution from orange to pink.

### PCR assay of Biopsy samples

The DNA template was extracted from each biopsy sample using HiPura™ mammalian genomic DNA purification Kit (Himedia, Mumbai, India).

### Amplification of 16S rDNA

The 16S rDNA (534 bp) of laboratory grown pure culture and DNA template of biopsy was amplified using *H. pylori*-specific primer^[10]^. The 16S rDNA gene primers of *H. pylori* were forward; 5′-TAAGAGATCAGCCTATGTCC-3′ and reverse primers; 5′-TCCCACGCTTTAAGCGCAAT-3′. Amplification was performed in a PTC 100 Thermal Cycler (MJ Research, Waltham, MA). The PCR reaction mix included 1.5 U of Taq DNA polymerase (Bangalore Genei, Bangalore, India), 1× PCR buffer with 1.5 mmol/l MgCl_2_ (1 mmol/l MgCl_2_ was added so as to attain a final concentration of 2.5 mmol/l), 25 pmol each of the forward and reverse primers (Integrated DNA Technologies, Coralville, IA), 125 μmol/l each of the dNTPs (deoxynucleotidetriphosphates), and 50 ng of template DNA or 3 µl suspension of obtained from biopsy in a total volume of 50 μl. Thermal cycles for the amplification were set at initial denaturation for 5 min at 94 °C, 40 cycles of 30 sat 94 °C, 30 s at 56 °C, and 1 min at 72 °C, followed by a final extension of 5 min at 72 °C. The amplified product (534 bp) was visualized on 1.2 % agarose gel using a gel documentation unit (BioRad Laboratories, Hercules, CA).

### Amplification of cagA gene

The cagA gene was amplified by using specific primer of cagA (350 bp) (including forward and reverse)^[11]^. The cagA gene primers were as follows: forward primer, 5′-GTTGATAACGCTGTCGCTTC-3’ and reverse primer, 5′-GGGTTGTATGATATTTTCCATAA-3’. The PCR mix included 0.9 U Taq DNA polymerase (Bangalore Genei), 1.5 mmol/l MgCl_2_ in standard PCR assay buffer (Bangalore Genei), 10 pmol each forward and reverse primer (Integrated DNA Technologies), 0.25 mmol/l each of the dNTPs and 50 ng template DNA or 3 μl of suspension (obtained from biopsy samples) in a total volume of 25 μl. The thermal program was set at initial denaturation for 3 min at 94 °C, 35 cycles of 1 min at 94 °C, 1 min at 55 °C, and 1 min at 72 °C, followed by the final extension of 10 min at 72 °C. Five microliters of the amplified PCR product was electrophoresed on a 2 % agarose gel containing 0.2 mg/ml ethidium bromide. The gels were run in Tris– borate–EDTA buffer and monitored as described above.

### Immunohistochemical Analysis

The methods of immunohistochemical analysis was used by the ToGA (Trastuzumab for gastric cancer) trial and also this method was adopted by the National Comprehensive Cancer Network guidelines version 2.201^[12,13]^. The procedure for collection of biopsy samples were slightly modified. The GC were fixed in 10 % neutral buffer formalin for 24 to 48 h and embedded in paraffin. The sections of 4 μm thickness were obtained from tissue blocks and processed for immunohistochemical (IHC) studies. Slides were then deparaffinised and rehydrated with descending grades of ethyl alcohol. IHC staining was done using the HER2 (BioGenex pre-diluted, Clone CB11, Mouse IgG) antibody. Immunostaining was carried out using a biotin–streptavidin complex. A 3, 3-Diaminobenzidinetetrahydrochloride was used as chromogen. Sections were counter stained with hematoxylin. In a microwave, slides were pre-treated with 10-mM citrated buffer, pH 6.0, at 95 °C, 5 min × 3 for antigen retrieval prior to incubation with primary antibody as above. Positive tissue controls of the gastric carcinoma as well as negative control slides that were run simultaneously and used to assess the quality of immunostaining. Immunoreactivity was quantified by evaluating a minimum of 1,000 carcinoma cells in randomly selected fields of histological sections on a slide, using Olympus Model B × 51 model. For the evaluation of HER2 expression, a score of 0 was given to sections showing no staining or membranous reactivity in < 10 % of cancer cells, and 1 + was given to sections showing faint or barely perceptible membrane reactivity in > 10 % of cancer cells; cells are reactive only in part of their membrane, 2 + was given to sections showing weak to moderate complete basolateral or lateral membranous reactivity in > 10 % of cancer cells, and 3+ was given to specimens showing strong complete basolateral or lateral membranous reactivity in > 10 % cancer cells. Scores of 0 and 1 + were considered negative for HER2 overexpression, whereas scores of 2 + and 3 + were considered to be HER2 overexpression.

### Amplification of HER2 gene

The amplification of *HER2* (154 bp) gene was performed by using *HER2* gene specific primer^[14]^. The primer sequence of the *HER2 gene* were forward: 5′-CCTCTGACGTCCATCGTCTC-3′ and reverse: 5′-CGGATCTTCTGCTGCCGTCG-3′. The PCR mix included 0.9 U Taq DNA polymerase (Bangalore Genei), 1.5 mmol/l MgCl_2_ in standard PCR assay buffer (Bangalore Genei), 10 pmol each forward and reverse primer (Integrated DNA Technologies), 0.25 mmol/l each of the dNTPs and 50 ng template DNA or 3 μl of suspension (obtained from biopsy samples) in a total volume of 25 μl. The thermal program was set at initial denaturation for 10 min at 95 °C, 40 cycles of 30 s at 94 °C, 1 min at 60 °C, and 1 min at 72 °C, followed by the final extension of 10 min at 98 °C. Five microliters of the amplified PCR product was electrophoresed on a 2 % agarose gel containing 0.2 mg/ml ethidium bromide. The gels were run in Tris–borate–EDTA buffer and monitored as described above.

### Interactive study of CagA

The 3D-structure of CagA (PDB ID: 4dvy)^[15]^ and HER2 (3pp0)^[16]^ were retrieved from protein data bank (PDB). Further the interactive potential of CagA for HER2 protein was analyzed by PatchDock server^[17]^. The web-based tool (http://anchor.enzim.hu) was implemented to predict the binding probability of CagA residues^[18]^.

### Statistical Analysis

The χ^2^ and Fisher exact tests were used for testing differences between the groups. The data are expressed as mean±standard deviation (S. D.). P values less than 0.05 and 0.01 were considered statistically significant. To further evaluate the correlation of HER2 status with CagA, we performed receiver operating characteristic (ROC) curve analysis by using DeLong et al. (1988) method. All quantitative data were statistically analyzed using MedCalc software version 12^[19]^.

## RESULTS

### Amplification of *H. pylori* specific genes in carcinoma tissues

PCR analysis was performed on 85 GC specimens. To confirm the presence of *H. pylori*, first RUT was performed with fresh biopsy samples. Of 85 specimens, RUT showed positive in 62 (72.94 %) biopsy samples and rest 23 biopsy showed negative test. Then, PCR assay was done to amplify the specific 16S rDNA of HP gene andamplified products were shown in the agarose gel photograph (Fig. 1a). Of 85 biopsies, 54 (63.52 %) showed excellent amplification for HP-specific 16S rDNA (534 bp) in above template recovered by HiPura™ kit and thirty-one specimens had not amplified (Fig. 1a). The incidence of risk by HP in both sex males and females were found significant (p = 0.039).

**Fig. 1.**
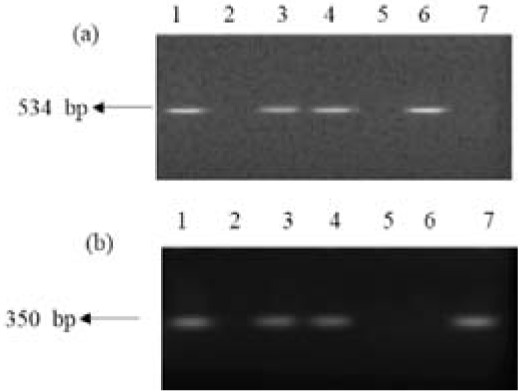
Representative PCR for amplification of 16S rDNA and cagA gene of *H. pylori* from biopsy of 5 cancerous patients, (a) for 16S rDNA- lane 1: ATCC 26695, lane 2: negative control, lane 3-7: different isolates, lane 5 and 7: no amplification. (b) for cagA gene- lane 1: ATCC 26695, lane 2: negative control, lane 3, 4, and 7 positive for cagA, lane: 5 and 6 no amplification.

Similarly, the HP virulent factor cagA (350 bp) in GC tissues was amplified by using specific primer of cagA gene and its product was shown in the agarose gel photograph (Fig. 1b). Of 54 biopsy of HP positive, 34 (62.96 %) GC tissues were amplified for cagA and 20 (37.03 %) specimens were negative. The frequency of cagA occurrence in HP infected tissues was insignificant (p = 0.3254).

### Detection of HER2 expression by Immunohistopathology and PCR amplification

The histology showed 57 (67.05 %) case diffuse type and 28 (32.94 %) intestinal type gastric cancer. Of 85 GC tissues, 17 male and 4 female specimens scored (2 + and 3 +) after immunostaining on IHC showed overexpression of HER2 and no overexpression (0 or 1 + immunostaining) in remaining 41 male and 23 female GC samples were observed [Fig. 2(a and b), Table 2a; P = 0.18]. In all, 28 patients had intestinal-type and 12 (42.85 %) of these were HER2 positive in comparison to 9 (15.79 %) of 57 GC tissues with diffuse-type cancer and showed statistically significant (Table 2a; P = 0.01).

**Table 2.** 2(a) Analysis of HER2 status in GC tissues by IHC.

| Criteria | HER2<br>positive (21) | HER2<br>negative (64) | p value |
| --- | --- | --- | --- |
| Gender |  |  | 0.18 |
| Male | 17 | 41 |  |
| Female | 4 | 23 |  |
| HPE |  |  | 0.01 |
| Diffuse | 9 | 48 |  |
| Intestinal | 12 | 16 |  |

$\chi^2$ test
HPE: Histopathological Examination

**Fig. 2.**
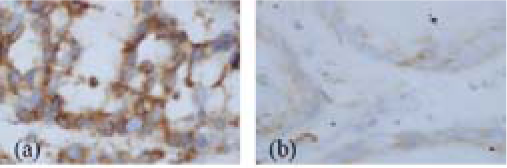
Immunoperoxidase staining of malignant cells for HER2 expression showing (a) maximum for positive (b) minimum for negative.

When we compared HER2 overexpression by IHC with HP status in GC tissues found in 17 (31.48 %) of 54 HP positive sample and 2 (0.065 %) of 31 HP negative sample. HER2 overexpression by IHC was closely associated with HP infection in gastric cancer (Table 2b P = 0.014). To further examine the association of *H. pylori* in expression of HER2 in biopsies, PCR for the HER2 gene was performed to confirm the presence of HER2 in the HP + isolates (Fig. 2c). A single amplicon was detected in 17 (20%) CagA + HP and 3 (5.55 %) in CagA - HP. The presence of CagA gene was closely associated with HER2 gene amplification (Table 2b, P = 0.0183).

2(b) Correlation between HER2 overexpression in *Helicobacter pylori* infected samples by immunohistochemistry and PCR amplification.

| Criteria | HER2+ | HER2- | P value |
| --- | --- | --- | --- |
| Overexpression by<br>IHC |  |  | 0.0141 |
| CagA+ HP | 15 | 19 |  |
| CagA- HP | 2 | 18 |  |
| Amplification by<br>PCR |  |  | 0.0183 |
| CagA+ HP | 17 | 17 |  |
| CagA- HP | 3 | 17 |  |

Fisher exact test
HP: *Helicobacter pylori*

The ability of the cagA gene to discriminate the HP with HER2 immunohistochemistry and amplification in *H. pylori* associated with biopsies from negative cases was also analysed by using ROC curve. The CagA predicted HER2 overexpression and amplification with an area under the ROC (AUC) values 0.750 and 0.875, respectively [Fig. 3(a) and (b)], suggesting that CagA may be a potential biomarker for HER2 status in gastric cancer.

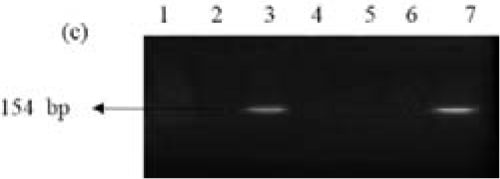
2(c) Representative results of *HER2* gene expression in different isolates with cagA+ and cagA- HP; lane 1: negative control, lane 2 and 5: no amplification of *HER2* gene in cagA+ HP isolate, lane 3: *HER2* gene amplification in cagA+ HP sample, lane 4 and 6: no amplification of *HER2* gene in cagA- HP isolate, lane 7: amplification of *HER2* gene in cagA- HP samples.

**Fig. 3.**
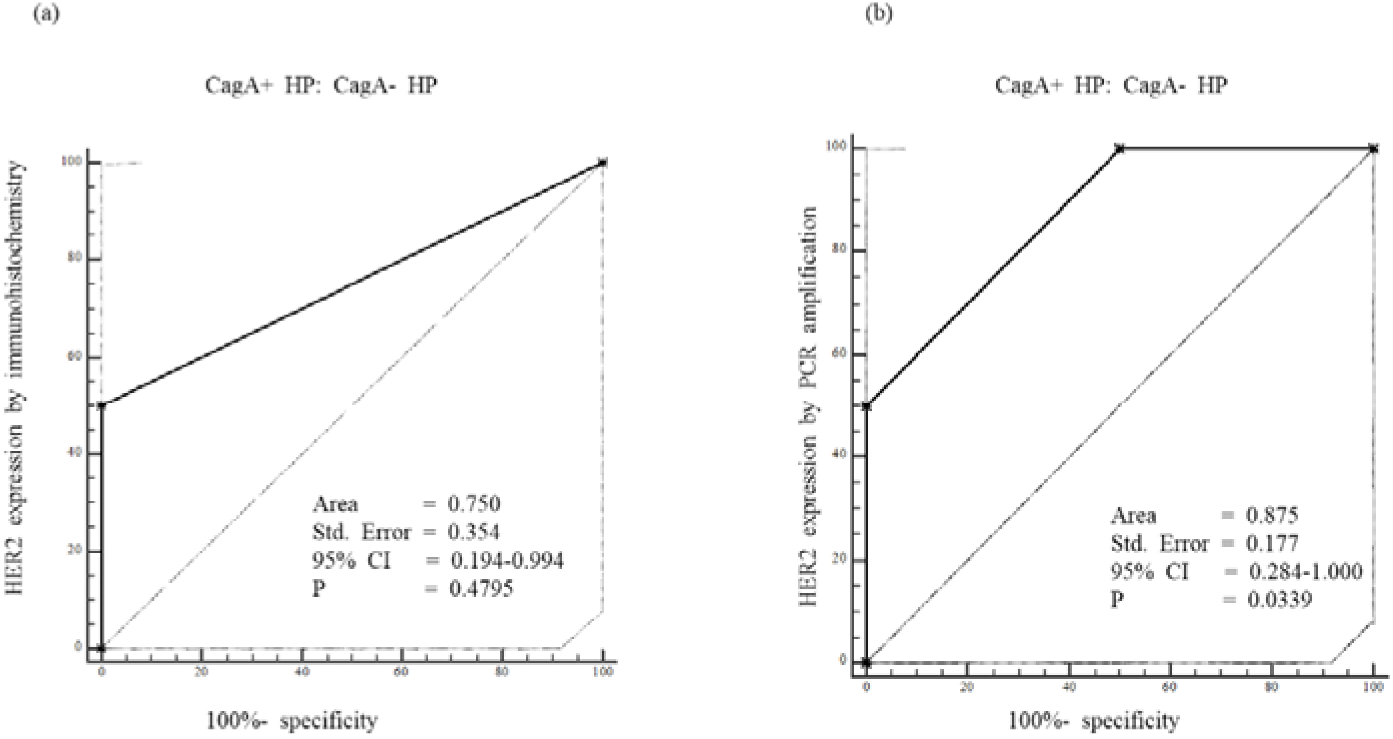
Association between the presence of CagA and HER2 status in *H. pylori* positive tissues. (a) The presence of CagA was closely associated with HER2 overexpression by IHC with an area under the ROC curve (AUC) value of 0.750 and (b) with HER2 amplification by PCR with an area under the ROC curve (AUC) value of 0.875.

### Interactions of CagA with HER2 protein

The molecular docking showed that the residues of CagA interacted to HER2 with binding energy −6.03 kcal/mol [Fig. 4(a)]. The blue ANCHOR graph of CagA revealed that the binding probability of CagA residues [Fig. 4(b)], which result was represented as percentage of binding probability of residues in scatter graph plot [Fig. 4(c)]. The scatter graph plot showed that the highest binding probability (84.6 %) in aromatic amino acid PHE864 and lowest (7.20 %) in hydroxyl containing THR862.

**Fig. 4.**
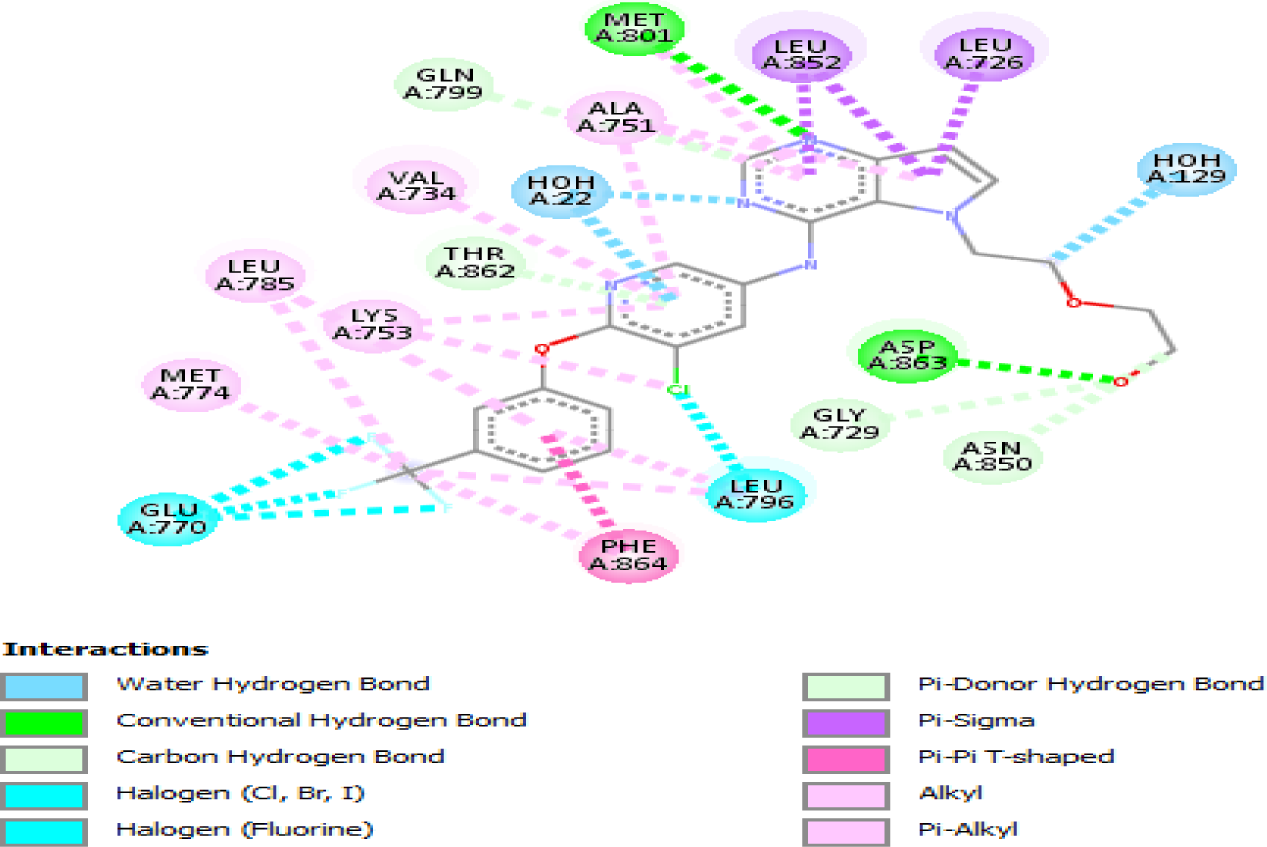
(a). Interactions of CagA residues to HER2 with binding energy (−6.03 kcal/mol)

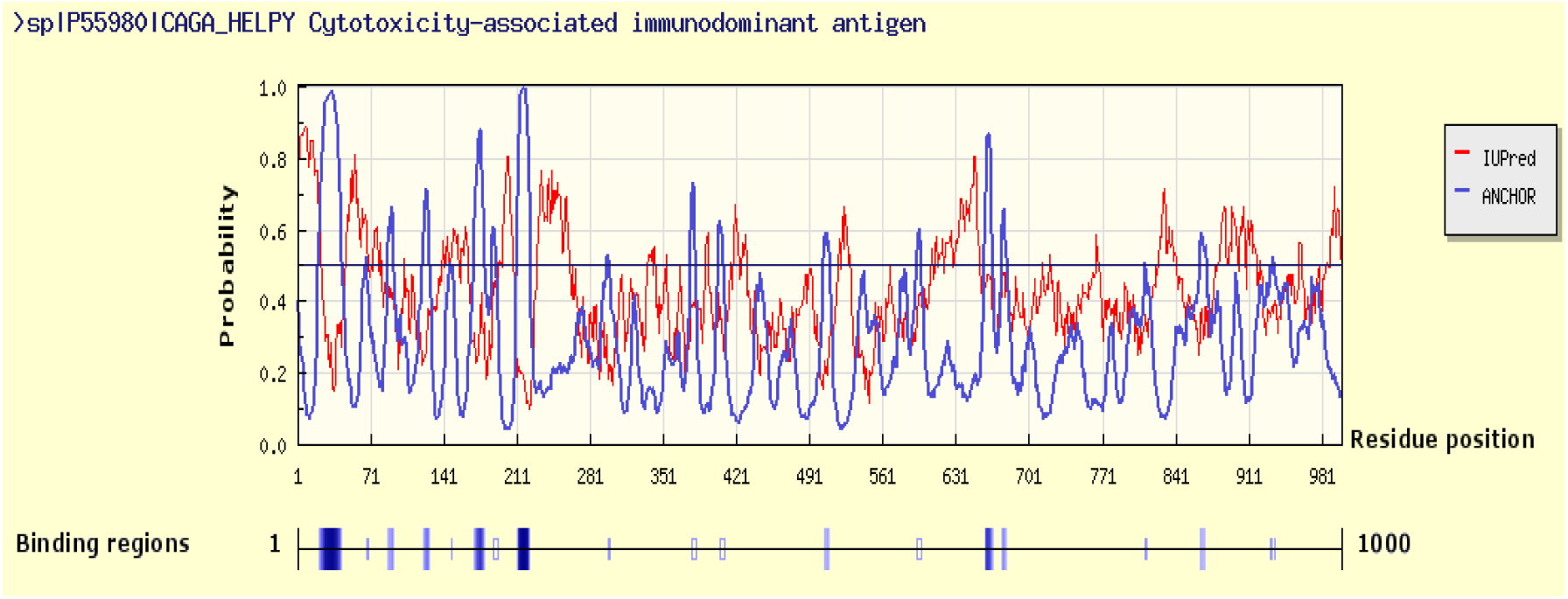
b. Anchor graph plot of CagA residues for prediction of its binding probability to HER2

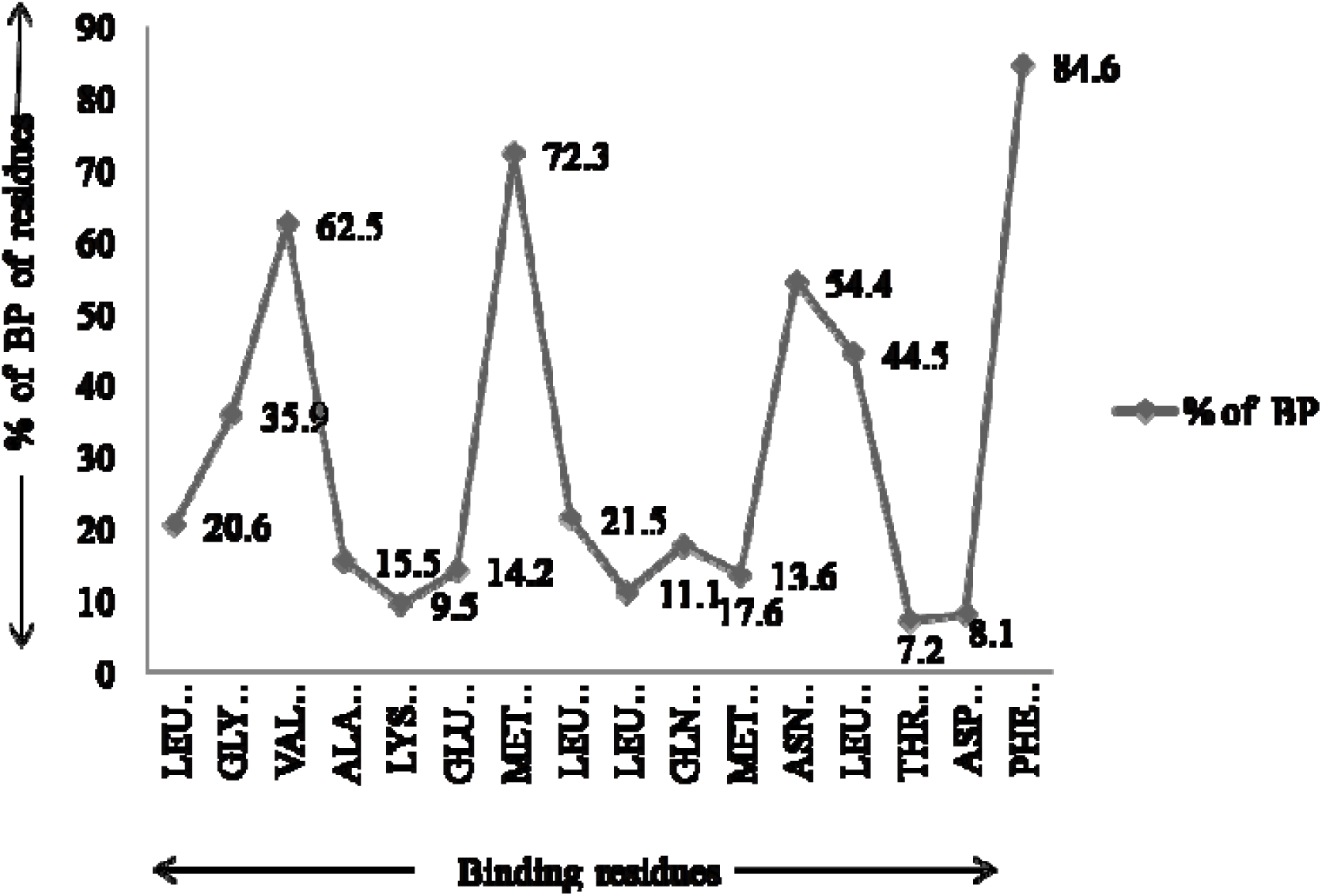
c. Scatter graph plot for presentation of binding probability (%) of CagA residues for HER2 expression.

## DISCUSSION

*H. pylori* has the capability to persist within hosts that recognizes its presence indicates important adaptations by changing environment, and involving programmed cross-interaction between microbe and host^[20]^. Kumar et al. (2008) suggested that the genotyping of *H. pylori* strains may be useful in clinical diagnosis of infected. patients^[21]^. The isolation of *H. pylori* from human biopsy by culture method is tedious process due to its longer incubation period required and fastidious growth nature. Hence, several alternative approaches have been generated for the accurate and rapid detection of *H. pylori* in gastric mucosa. The genomic diversity among *H. pylori* after clinical isolation has been assayed using 16S rDNA sequence and numerous sequence-specific PCR assays to identify *Helicobacter* species^[22]^. Therefore, 16s rDNA-based PCR methods have been adopted for rapid detection of non-culturable bacteria in specimens of human biopsy^[23]^. 16S rDNA gene ordinarily exists in the genome of *H. pylori.* However, its sensitivity is determined by the primers selected for *H. pylori* specific 16S rDNA, the source of DNA and quantity of bacterial presented in cancerous tissues. A typical PCR amplification of the 16S rDNA gene is shown in Fig. 1(a). The size of the amplified nucleotide fragment was 534 bp similar to used *H. pylori* specific gene 16S rDNA and this alternative approach by PCR assay made confirm to be presence of *H. pylori* in cancerous tissues. Of 85 patients, fifty four (62.96 %) were positive for HP infection, and these patients were designated as tissue positive. Other 31 biopsies of gastric cancer were not amplified of the desired 16S rDNA gene and showed the tissues negative for HP. The small size of gastric biopsy specimen or minor colonisation of pathogens may results of colonised and non-colonised tissues^[24]^.

An important virulence factor cagA gene of *H. pylori* associated in the development of cancer, was found to be common irrespective of the clinical diagnosis, similar to previous study by Smith et al. (2002)^[25]^. Furthermore, total genomic DNA of 54 HP positive samples were used for amplification of nucleotide fragment cagA gene using specific primer. Fig. 1(b) showed agarose gel photograph of amplified product for cagA (350 bp) gene. Out of 54 HP positive samples, 34 (62.96 %) were positive for cagA, and remaining 20 (37.03 %) specimens were negative. The results indicated thatthe distribution of virulent factor cagA gene were not same in all *H. pylori* strain. As a study have explained that the cagA is not found in all *H. pylori* strains^[26]^.

Epidermal growth factor receptors (EGFR) family have four genes coding of four homologous epidermal growth factor receptors^[27]^. All receptors are localised in cellular membrane of various tissues and code for transmembrane glycoprotein referred as HER-2 (human epidermal growth factor receptor-2) protein or receptor. HER2 overexpression and amplification occurs in several cancers including gastric cancer^[28,29]^. *H. pylori* plays a vital role in expression of EGF receptor in gastric tissues and that leads to growth, proliferation, and differentiation in tissues^[30]^. This overexpression can cause of gastric cancer development and ended up with HER2 oncogene activities^[30]^. This is one of spontaneous mechanisms involving in changing of normal cellular activities for development of gastric cancer. HER2 status in GC patients was observed by IHC and PCR method. The maximum immunostaining for positive and minimum staining for negative expression of HER2 were shown in Fig. (2). In this series, out of 85 biopsies of gastric adenocarcinoma, HER2 overexpression was observed in 21 (24.70 %) cases on IHC and 64 (75.29 %) biopsy of gastric cancer showed negative expression (Table 2a). Moelans et al. (2011) have reported that the positive expression of HER2 protein is minimum in patients of gastric cancer ranging from 2-45%^[31]^. Histopathology results also showed that intestinal-type cancers showed higher (75 %) HER2 expression than diffuse type (18.75 %) which supported to previous findings of Park et al. (2006)^[32]^.

Similarly, we investigated the above obtained results of HER2 expression with cagA+ *H. pylori* in carcinoma tissues to reveal the relations and represented as Fig 3(a, b, and c). We examined and compared the presence of CagA with HER2 status in gastric tissues. Interestingly, a positive correlation was observed between HER2 overexpression and amplification with CagA positivity (Table 2b; P = 0.0141, P = 0.0183). When the ability of CagA to discriminate the gastric cancer with HER2 expression and amplification was analysed using an ROC curve, CagA predicted HER2 amplification and overexpression in tissues with AUC values of 0.750 and 0.875, respectively [Fig. 3(a and b)]. Finally, we conclude that CagA may be one of the candidate biomarker for HER2 amplification and contribute to the development of gastric cancer by inducing HER2 amplification and overexpression. Shim et al. (2014) have also reported that CagA could be considered as biomarker for HER2 overexpression or amplification in gastric cancer^[33]^. Similarly, the relation of HER2 overexpression and hormones receptors status in breast cancer has been established^[34]^.

The molecular docking showed that the HER2 was sensitized to interact with residues of CagA with binding energy −6.03 kcal/mol [Fig. 4(a)]. The ANCHOR graph of CagA revealed that the binding probability of CagA residues measured its disorderness [Fig. 4(b)] that could be disrupt the natural integrity of HER2. The highest binding probability (84.6 %) in aromatic amino acid PHE864 and lowest (7.20 %) in hydroxyl containing THR862 [Fig. 4(c)] may not change HER2 confirmation uniformly.

This study concluded that cagA+ *H. pylori* strain in human may damage the activities of host proteins and nucleotides causing more prone to gastric cancer. It has been investigated that the CagA+ *H. pylori* strain may induce for HER2 expression in gastric cancer. The molecular docking study also showed that the CagA could disrupt the natural integrity of HER2 proteins, which may express in cancerous tissues. The emergence cagA in tissues associated with gastric cancer and specific reasons of HER2 overexpression in this cancer are very complex and need further more study to improve quality in diagnosis as well as therapy for human kinds.

## ACKNOWLEDGMENT

The authors are thankful to University Grants Commission for financial support. The authors are also thankful to Prof. B.K. Roy, Department of Botany, Banaras Hindu University, India for revising manuscript.

## CONFLICT OF INTEREST

The authors declare no conflicts of interest.

